# COVFlow: phylodynamics analyses of viruses from selected SARS-CoV-2 genome sequences

**DOI:** 10.1101/2022.06.17.496544

**Authors:** Gonché Danesh, Corentin Boennec, Laura Verdurme, Mathilde Roussel, Sabine Trombert-Paolantoni, Benoit Visseaux, Stéphanie Haim-Boukobza, Samuel Alizon

**Affiliations:** MIVEGEC, CNRS, IRD, Université de Montpellier; Laboratoire CERBA, Saint Ouen L’Aumône, France; Center for Interdisciplinary Research in Biology (CIRB), College de France, CNRS, INSERM, Université PSL, Paris, France

**Keywords:** COVID-19, molecular epidemiology, sequence database, phylogenetics, public health

## Abstract

Phylodynamic analyses can generate important and timely data to optimise public health response to SARS-CoV-2 outbreaks and epidemics. However, their implementation is hampered by the massive amount of sequence data and the difficulty to parameterise dedicated software packages. We introduce the `COVFlow` pipeline, accessible at https://gitlab.in2p3.fr/ete/CoV-flow, which allows a user to select sequences from the Global Initiative on Sharing Avian Influenza Data (GISAID) database according to user-specified criteria, to perform basic phylogenetic analyses, and to produce an XML file to be run in the `Beast2` software package. We illustrate the potential of this tool by studying two sets of sequences from the Delta variant in two French regions. This pipeline can facilitate the use of virus sequence data at the local level, for instance, to track the dynamics of a particular lineage or variant in a region of interest.

## 1 Introduction

Millions of SARS-CoV-2 full genome sequences have been generated since 2020, and, for the majority, been made available through the Global Initiative on Sharing Avian Influenza Data (GISAID) consortium [1, 2]. This has allowed the timely monitoring of variants of concerns (VoC) with platforms such as CoVariants (CoVariants), outbreak.info [3], or CoV-Spectrum [4], and the realisation of phylogenetic analyses, e.g. via NextStrain [5].

Phylogenies represent a powerful means to analyse epidemics with an intuitive parallel between a transmission chain and a time-scaled phylogeny of infections, which is the essence of the field known as ‘phylodynamics’ [6]. As illustrated in the case of the COVID-19 pandemic, state-of-the-art analyses allow one to investigate the spatio-temporal spread of an epidemic [7], superspreading events [8], and even detect differences in transmission rates between variants [9].

Phylodynamic analyses involve several technical steps to go from time-stamped virus sequence data to epidemiological parameter estimates, which can make them difficult to access to a large audience. Furthermore, the amount of data shared greatly overcomes the capacities of most software packages and imposes additional selection steps that further decrease the accessibility of these approaches. To address these limitations, we introduce the `COVflow` pipeline which covers all the steps from filtering the sequence data according to criteria of interest (e.g. sampling data, sampling location, virus lineage, or sequence quality) to generating a time-scaled phylogeny and an XML configuration file for a BDSKY model [10] to be run in the Beast2 software package [11].

Some pipelines already exist to assess sequence quality, filter data, infer an alignment, and infer a time-scaled phylogeny such as Nextclade [12] and Augur [13]. However, these do not include a step to perform a phylodynamic analysis from the output files, which requires dedicated skills. The `COVFlow` pipeline addresses this limitation and integrates all the steps present in separate software packages to go from the sequence data and metadata to the XML to be run in Beast2.

Here, we present the architecture of the pipeline and apply it to data from the French epidemic, which has been poorly analysed (but see [14–16]). Focusing on sequences belonging to the Delta variant collected in France in two regions, Ile-de-France, and Provence-Alpes-Cote-d’Azur, by a specific French laboratory (CERBA), we illustrate the pipeline accessibility, flexibility, and public health relevance.

## 2 Methods

`COVFlow` is a bioinformatics pipeline for phylogenetic and phylodynamic analysis of SARS-CoV-2 genome sequences. It is based on the Snakemake workflow management system [17] and its dependencies are easily installed via a conda virtual environment. Snakemake ensures reproducibility, while Conda (https://docs.conda.io/en/latest/) and Bioconda [18] allows for version control of the programs used in the pipeline. Overall, the pipeline is easy to install and avoids dependency conflicts.

### Pipeline configuration

The pipeline workflow is configured using a YAML configuration file, which must contain the path to the sequence data file, the path to the metadata file, and the prefix chosen for the output files. Each parameter of the pipeline following steps has a default value, which can be modified by the user in the configuration file.

### Input data

The input data analysed by `COVFlow` are sequence data and metadata, corresponding to patient properties, that can be downloaded from the Global Initiative on Sharing All Influenza Data (GISAID, https://www.gisaid.org/). The sequence data are in a FASTA format file. The metadata downloaded contains details regarding the patient’s sequence ID (column named ‘strain’), the sampling dates (column ‘date’), the region, country, and division where the sampling was made (columns respectively named ‘region’, ‘country’, and ‘division’). It also lists the virus lineage assigned by the Pangolin tool [19], and the age and sex of the patient (columns respectively named ‘pango_lineage’, ‘age’, and ‘sex’).

### Data filtering

The first step implemented in the pipeline performs quality filtering. By default, genomic sequences that are shorter than 27, 000 bp, or that have more than 3, 000 missing data (i.e. N bases) and more than 15 non-ATGCN bases are excluded. These parameter values can be modified by the user. Sequences belonging to non-human or unknown hosts are also excluded. Sequences for which the sampling date is more recent than the submission date, or for which the sampling date is unclear (e.g. missing day) are also excluded. Finally, duplicated sequences and sequences that are flagged by the Nextclade tool [12] with an overall bad quality (Nextclade QC overall status ‘bad’ or ‘mediocre’) are also removed.

The sequence data is then further filtered following the user’s criteria. These include Pangolin lineages, sampling locations (regions, countries, or divisions), and sampling dates. In addition to specifying the maximum and/or minimum sampling dates, the user can specify a sub-sampling scheme of the data with a number or percentage of the data per location and/or per month. For example, the user can decide to keep *x*% of the data per country per division per month or to keep *y* sequence data per division. Finally, more specific constraints can be given using a JSON format file with three possible actions: i) keep only rows (i.e. sequences) that match or contain a certain value, ii) remove rows that match or contain a certain value, and iii) replace the value of a column by another value for specific rows with a column that matches or contains a certain value. The last action can be used to correct the metadata, for instance, if the division field is not filled in but can be inferred from the names of the submitting laboratory. The JSON file can be composed of multiple key-value pairs, each belonging to one of the three actions. For example, the user can specify to keep only male patients and to remove data from one particular division while setting the division of all the samples submitted by a public hospital from the Paris area (i.e. the APHP) to the value ‘Ile-de-France’.

### Aligning and masking

The set of sequences resulting from the data filtering is then divided into temporary FASTA files with a maximum number of 200 sequences per file. For each subset, sequences are aligned to the reference genome MN908947.3 using `MAFFT` v7.305 [20] with the ‘keeplength’ and ‘add’ options. All the aligned sequences are then aggregated into a single file. Following earlier studies, the first 55, the last 100 sites, and other sites recommended from https://github.com/W-L/ProblematicSites_SARS-CoV2 of the alignment are then masked to improve phylogenetic inference (http://virological.org/t/issues-with-sars-cov-2-sequencing-data/473).

### Inferring and time-scaling a phylogeny

A maximum-likelihood phylogenetic tree is estimated using `IQ-TREE` v2.1.2 [21] under a GTR substitution model from the alignment. The resulting phylogeny is time-scaled using `TreeTime` v0.8.1 [22]. By default, the tree is rooted using two ancestral sequences (Genbank accession numbers MN908947.3 and MT019529.1) as an outgroup, which is then removed, with a fixed clock rate of 8 · 10^−4^ substitutions per position per year [23] and a standard deviation of the given clock rate of 0.0004. These parameters can be modified by the user. The output phylogeny is in a Newick format file.

### BDSKY XML file generating for BEAST 2

The Bayesian birth-death skyline plot (or BDSKY) method allows the inference of the effective reproduction number from genetic data or directly from a phylogenetic tree, by estimating transmission, recovery, and sampling rates [10]. This method allows these parameters to vary through time and is implemented within the `BEAST 2` software package [11].

Performing a BDSKY analysis requires setting an XML file specifying the parameters for the priors. As in any Bayesian analysis, this step is extremely important. The default settings in BEAST2 have been chosen to minimise the risk of errors. `COVFlow` builds on most of these with some modifications to fit the needs of large SARS-CoV-2 phylogenies.

The most important change has to do with the inference of the phylogeny. This can be done by `BEAST 2` but to minimise computation speed and allow for the analysis of large phylogenies, the pipeline sets the time-scaled phylogeny from the previous step in the XML file.

The default XML file assumes that there are two varying effective reproduction numbers to estimate, with a lognormal prior distribution, LogNorm(*M* = 0, *S* = 1), resulting in a median of 1, the 95% quantiles falling between 7.10 and 0.14, and a starting value of 1.0. This prior is adapted to such virus epidemics and, as we will see below, can be edited if needed. The default prior for the rate of end of the infectious period is a uniform distribution, Uniform(10, 300), resulting in a median of 155[17.3; 293]years^−1^, with a starting value of 70 years^−1^, and is assumed to be constant over time. The inverse of the rate of end of the infectious period is the average infectious period. This default prior yields infectious periods varying from 0.034 year (1.2 days) to 0.1 year (36.5 days), which is consistent with the biology of SARS-CoV-2 infections [24]. Usually, little or no sampling effort is made before the first sample was collected. Therefore, by default, we assume two sampling proportions: before the first sampling date it is set to zero, and after the default prior is a beta distribution, Beta(*α* = 1, *β* = 1), with a starting value of 0.01, translating in a median of 0.50 ([0.025; 0.975]). The non-zero sampling proportion is assumed to remain constant during the time the samples were collected. The method can also estimate the date of origin of the index case, which corresponds to the total duration of the epidemic. Since the tree is assumed to be a sampled tree, and not a complete one, the origin is always earlier than the time to the most recent common ancestor of the tree. Hence, the prior distribution’s starting value and upper value must be higher than the tree height. This condition is always checked when running the pipeline. The default prior for this parameter prior is a uniform distribution Uniform(0, height + 2) years, with a starting value of height, with height as the maximum height of the inferred time-scaled tree.

Note that, although the default priors are designed to minimise the risk of bias in the results and the pipeline checks for the origin parameter prior, the choice of the priors is essential and may impact the phylodynamic inference of parameters.

In the `COVFlow` configuration file, the user can modify the distribution shapes, the starting values, the upper and lower values, and the dimensions for each of these parameters to estimate, and set the dates at which the parameter estimation changes. The length of the MCMC chain and the sampling frequency, which are by default set to 10, 000, 000 and 100, 000 respectively, can also be modified.

The `BEAST 2` inference itself is not included in the pipeline. The reason for this is that a preliminary step (i.e. installing the BDSKY package) needs to be performed by the user. Similarly, the analysis of the BEAST2 output log files needs to be performed by the user via Tracer [25] or a dedicated R script available on the `COVflow` Gitlab page.

Compared with the `Nextclade` pipeline [5], `COVflow` allows a more flexible filtering stage using the JSON file. For example, it can select data if a column contains a certain word, allowing the user to filter data that may contain spelling mistakes or to select data from a group of laboratories that contain a common word (in our case CERBA) but don’t have the same names. Furthermore, the sub-sampling can either be based on the number of data points or on the percentage of available data and the latter option is currently not possible with `Nextstrain`. The masking sites strategy is also different between the two pipelines. Finally, and perhaps most importantly, `COVflow` configures an XML file for a BDSKY phylodynamic analysis in Beast 2, allowing for more detailed phylodynamic analyses.

### Illustration study with French data

We applied `COVFlow` to analyse GISAID data by downloading sequence data and metadata from the GISAID platform for the GK clade corresponding to the lineage B.1.617.2 available on April 22, 2022, which amounted to 4, 212, 049 sequences. Using the pipeline and the editing of its JSON file, we cleaned the sequence data, selected the data collected by a specific large French laboratory (CERBA), selected the data from two regions of interest (the Ile-de-France region for a first analysis, and the Provence-Alpes-Côte d’Azur region for a second analysis), and sub-sampled the data to keep up to 50 sequences per month. These two regions were chosen because they had some of the highest coverage in the dataset, while being in different parts of France. Our third analysis included the whole country so we sub-sampled the data to keep up to 50 sequences per month per French region. The other parameters of the pipeline were default except for the number of windows for the effective reproduction numbers in the BDSKY analysis which was set to 9 with a change-point time every month from June 01, 2021, to January 01, 2022.

To evaluate the robustness of the inference, we performed 5 independent `COVFlow` runs for France, using the pipeline configuration described above for the France analysis. For each run, we manually extracted the two major clades representing at least 20% of the leaves from the resulting phylogeny and used a Python script of the `COVFlow` pipeline to generate two XML files. For each BDSKY analysis, 9 effective reproduction numbers were estimated over the same time periods.

To assess the validity of the BDSKY results, we extracted SARS-CoV-2 PCR screening data from https://www.data.gouv.fr/fr/datasets/r/5c4e1452-3850-4b59-b11c-3dd51d7fb8b5. More precisely, we used the positivity rate at the national level and in the two regions of interest. The effective reproduction number (*R*_*e*_) was estimated using the `EpiEstim` R package [26, 27]. The data were smoothed out using a 7-days rolling average, in order to compensate for the reporting delays.

The files necessary to generate these results are provided in Appendix.

## 3 Results

We illustrate the potential of the `COVFlow` pipeline by performing a phylodynamic analysis of a specific COVID-19 lineage, here the Delta variant (Pango lineage B.1.617.2), in two regions of a country, here Ile-de-France and Provence-Alpes-Côte d’Azur in France (Figure 2(a)).

**Figure 1:**
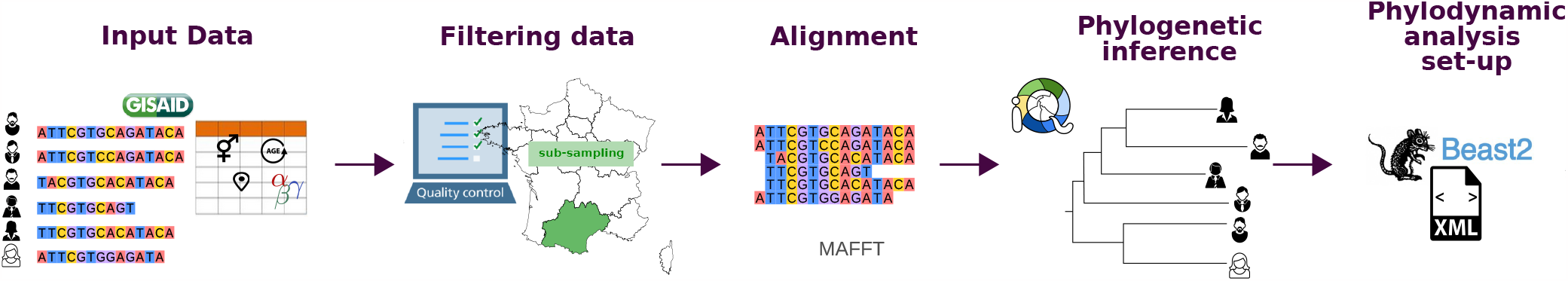
Structure of the COVFlow pipeline. The input data correspond to FASTA sequences and metadata provided by the GISAID. The data filtering is done using a YAML configuration file. The sequence alignment is performed with `MAFFT` and the phylogenetic inference with `IQ-TREE`. The pipeline generates an XML file that can be directly used with `Beast2`.

**Figure 2:**
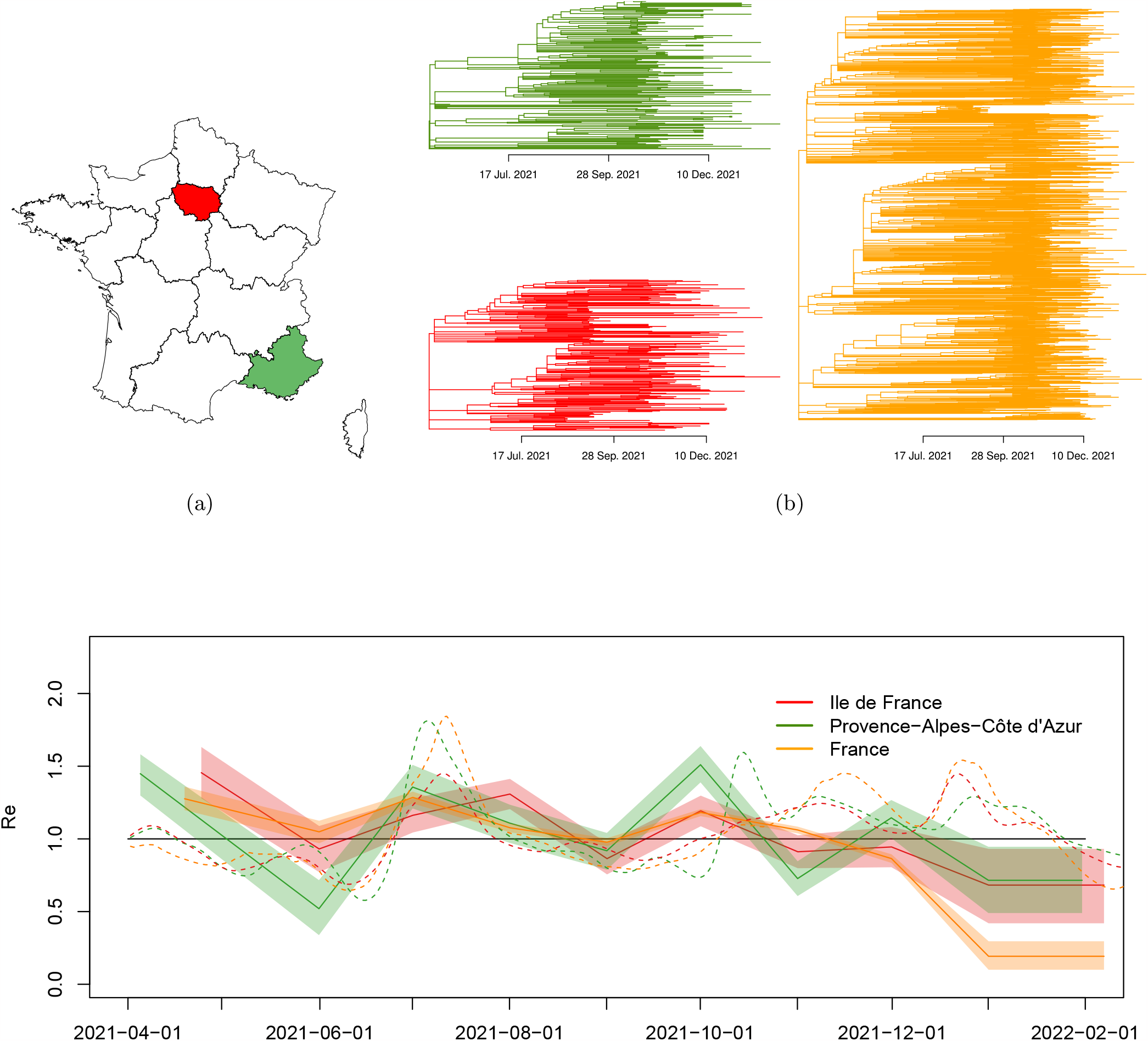
Analysing the SARS-CoV-2 Delta variant epidemics in French regions using the COVFlow pipeline. a) Geographical sub-sampling using at most 50 sequences per month for the Delta variant in Ile-de-France (IdF, in red), Provence-Alpes-Côte d’Azur (PACA, in green), and in all of France collected by CERBA laboratory. b) Time-scaled phylogenies generated using sub-sampled data from IdF (in red), PACA (green), and all of France (in orange). c) Temporal variations of the effective reproduction number (*R*_*e*_) of the Delta variant in IdF (red), in PACA (green), and France (orange) estimated using Beast2 from phylogenies in solid lines, and estimated using Epiestim from incidence data in dashed lines. The last panel was generated using Beast2. In panel c, the solid lines show the median values and the shaded area the 95% highest posterior density.

The `COVflow` runs resulted in the selection of 176 SARS-CoV-2 genomes for Ile-de-France (IdF), 221 genomes for Provence-Alpes-Côte d’Azur (PACA), and 1, 575 genomes for France.

The first output of the pipeline is the time-scaled phylogeny inferred from the sequences. In Figure 2(b), we show the one for each of the two regions considered and the one for the whole country. This already allows us to visualise the date of origin of the epidemic associated with the sequences sampled. More generally, the shape of the phylogenies can reflect the epidemic spread in the locality studied, e.g. the number of external introductions.

The second output of the pipeline is the XML file for a BDSKY model that can be run into Beast2. In Figure 2(c), we show the temporal variations in the effective reproduction number (*R*_*e*_), which is the average number of secondary infections caused by an infected individual at a given date. If *R*_*e*_ *<* 1, the epidemic is decreasing and if *R*_*e*_ *>* 1 it is growing.

The results show that the Delta variant epidemic seems to have started earlier in PACA than in IdF in early 2021. In both regions (and in France), the growth of the Delta variant in June is consistent with previous results showing the transmission advantage of 79% over the Alpha variant during this time period [28]. Furthermore, the earlier start in PACA is consistent with the beginning of the school holidays, PACA being a densely populated region in the summer. Note that IdF, as PACA, was more above the French average, which is also unsurprising given the density and international connections of the region.

Early in the fall of 2021, the back-to-school period led to an epidemic rebound in France. The associated epidemic growth was again stronger in PACA than the national average. Furthermore, contrarily to IdF or France, PACA experienced a period of Delta variant growth following the winter holidays. These are more difficult to explain but could be linked to local differences in terms of behaviour.

Finally, we see a clear slowdown in the Delta variant epidemic at the end of 2021. This is likely linked to the extension of the 3^rd^ vaccination dose, to changes in French behavior, but also to emergence of Omicron BA.1 variant, which was shown to have a growth advantage over the Delta variant [29].

When comparing BDSKY estimates with that of `EpiEstim` on the screening tests (dashed lines), we generally found consistent results. However, we did observe a shift in *R*_*e*_ peaks. This is consistent with the fact that methods based on incidence have an intrinsic delay due to the lag between the date of the infection and that of the PCR testing. For phylodynamics, this delay is, in theory, less important since the methods focus on virus evolution. `EpiEstim` estimates detect an epidemic growth in IdF and then PACA at the end of 2021 but this is expected because PCR tests do not discriminate between lineages and the end of the 2021 year saw the rise of the Omicron BA.1 variant [29].

Finally, a worry with phylodynamics is that the results depend on the sequences chosen. Moreover, considering the whole phylogeny incorporates importation events that are not included explicitly in the underlying birth-death model assumed by the BDSKY methods. Fig. 3, we show that the effective reproduction numbers estimated from the main subtrees are quantitatively similar to the *R*_*e*_ estimated from the whole phylogenetic tree. Furthermore, the estimations are all similar for different runs suggesting that the BDSKY framework is robust to phylogenetic tree uncertainty.

**Figure 3:**
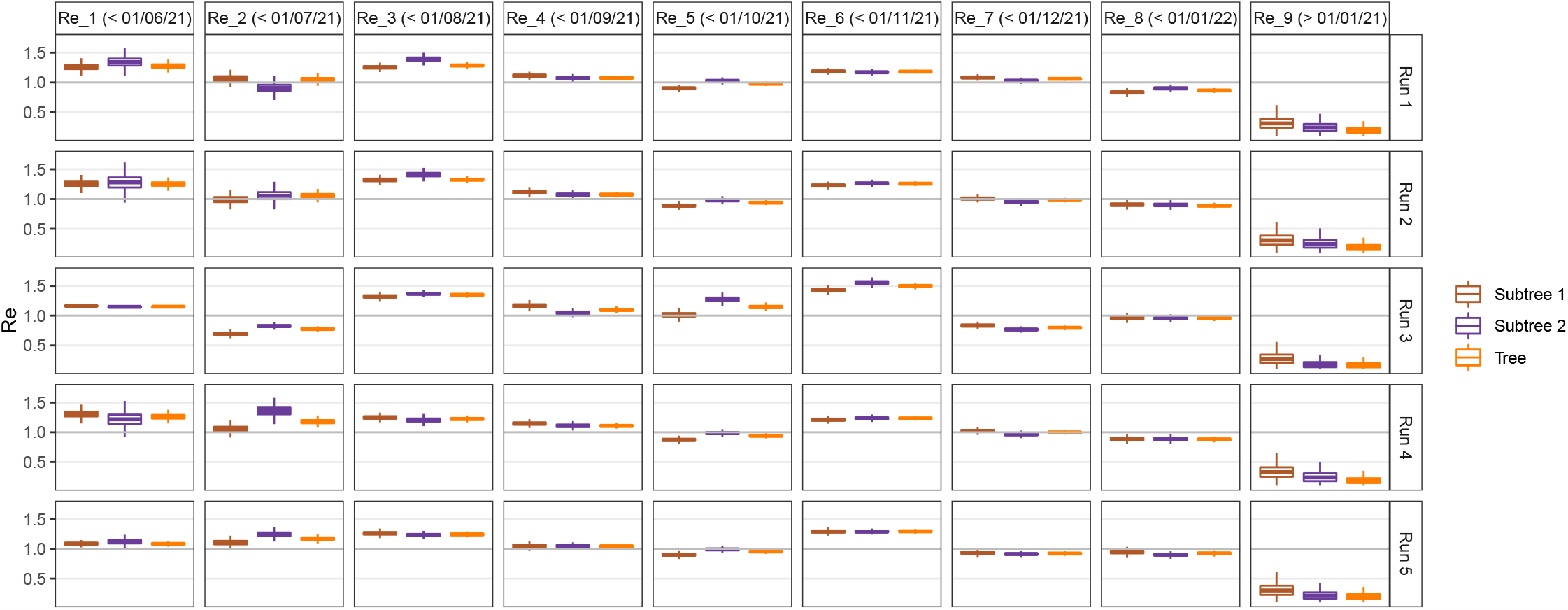
Estimations of the effective reproduction number *R*_*e*_ of the Delta variant in France for 5 different COVflow runs. For each run, the *R*_*e*_ were estimated from the inferred phylogenetic tree, and for the two principal clades, denoted Subtree 1 and Subtree 2. For each tree, 9 different *R*_*e*_ were estimated, with a changing point date every month from 2021-06-01 to 2022-01-01.

## 4 Discussion

The COVID-19 pandemic constitutes a qualitative shift in terms of the generation, sharing, and analysis of virus genomic sequence data. The GISAID initiative allowed the rapid sharing of SARS-CoV-2 sequence data, which is instrumental for local, national, and international public health structures that need to provide timely reports on the sanitary situation. At a more fundamental level, this genomic data is also key to furthering our understanding of the spread and evolution of the COVID-19 pandemic [30], especially in low-resource countries [31].

We elaborated the `COV-flow` pipeline, which allows users to perform all the steps from sequence data to phylodynamics analyses. In particular, it can select sequences from the GISAID dataset based on metadata, perform a quality check, align the sequences, infer a phylogeny, root this phylogeny into time, and generate an XML file for `Beast2` analysis (we also provide scripts to analyse the outputs). Furthermore, `COV-flow` can also readily allow the implementation of subsampling schemes per location and per date. This can help balance the dataset and also be extremely useful to perform sensitivity analyses and explore the robustness of the phylodynamic results.

A future extension could consist in including other `Beast2` population dynamics models, for instance, the Bayesian Skyline model, which is not informative about *R*_0_ but is potentially less sensitive to variations in sampling intensity. Another extension could be to use other databases to import SARS-CoV-2 genome data, e.g. that published by NCBI, via LAPIS (Lightweight API for Sequences).

Beast2 can simultaneously infer population dynamics parameters and phylogenies, which is an accurate way to factor in phylogenetic uncertainty [11]. However, this global inference is particularly computationally heavy and is out of reach for large data sets. To circumvent this problem, we perform the phylogenetic inference first using less accurate software packages and then impose the resulting phylogeny into the Beast2 XML file. An extension of the pipeline could offer the user to also perform the phylogenetic inference, for instance by using the so-called ‘Thorney Beast’ (https://beast.community/thorney_beast) implemented in Beast 1.10 [32].

Finally, it is important to stress that phylogenetic analyses are always dependent on the sampling scheme [33–36]. If most of the sequences come from contact tracing in dense clusters, the analysis will tend to overestimate epidemic spread. This potential bias can be amplified by the sequence selection feature introduced in the pipeline. An advantage of `COVFlow` is that it can perform spatio-temporal subsampling but additional studies are needed to identify which are the most appropriate subsampling schemes to implement.

## Acknowledgement

The authors acknowledge further support from the CNRS, the IRD and the i-Trop HPC (South Green Platform) at IRD Montpellier, which provided HPC resources that contributed to the results reported here (https://bioinfo.ird.fr/).

The authors thank the Experimental and Theoretical Evolution team from Maladies Infectieuses et Vecteurs: Écologie, Génétique, Évolution et Contrôle, University of Montpellier, for discussion, as well as the EMERGEN consortium (complete member list in Supplementary Materials).

This project was supported by the Agence Nationale de la Recherche Maladies Infectieuses Émergentes to the MODVAR project (grant no. ANRS0151).

## Authors contributions

GD and SA conceived the study, GD built the pipeline and performed the analyses, CB contributed to the implementation of the pipeline, LV, MR, STP, BV, and SHB contributed genetic sequence data, SA and GD wrote a first version of the manuscript.

## Data and scripts

The sequences analysed were generated by CERBA and uploaded to GISAID.

The R scripts, along with all the files generated by the pipeline and used for the analyses (XML files, FASTA alignments, time-scaled phylogenies) are provided in Supplementary Materials.

The pipeline itself can be accessed on the Git public repository https://gitlab.in2p3.fr/ete/CoV-flow

## Conflict of Interest

The authors of this preprint declare that they have no financial conflict of interest with the content of this article.

## Notes

https://zenodo.org/record/8016623

### Competing Interest Statement

The authors have declared no competing interest.

### Summary of Updates

Title changed

